# Inhibiting circRNA *Cdr1as* expression in the ILPFC of adult male C57BL/6J mice impairs fear extinction memory

**DOI:** 10.1101/2022.07.14.500137

**Authors:** Esmi Lau Zajaczkowski, Qiongyi Zhao, Wei-Siang Liau, Hao Gong, Sachithrani Umanda Madugalle, Ambika Periyakaruppiah, Laura Jane Leighton, Mason Musgrove, Haobin Ren, Joshua Davies, Paul Robert Marshall, Timothy William Bredy

**Author notes:** Co-corresponding authors: Paul Robert Marshall and Timothy William Bredy.

## Abstract

Circular RNAs (circRNAs) comprise a novel class of regulatory RNAs that are abundant in the brain, particularly within synapses. They are highly stable, dynamically regulated, and display a range of functional roles, including as decoys for miRNAs and proteins and, in some cases, translation. Early work in animal models revealed an association between circRNAs and neurodegenerative and neuropsychiatric disorders; however, relatively few studies have shown a causal link between circRNA function and memory. To address this knowledge gap, we sequenced circRNAs in the synaptosome compartment of the medial prefrontal cortex of fear extinction trained male C57BL/6J mice and found 12837 circRNAs enriched at the synapse, including *Cdr1as*. Targeted knockdown of *Cdr1as* in the neural processes of the infralimbic prefrontal cortex of male C57BL/6J mice led to impaired fear extinction memory. Altogether, our findings highlight the importance of localised circRNA activity at the synapse for memory formation and suggest that circRNAs may have a more widespread effect on brain function than previously thought.

## 1. Introduction

Circular RNAs (circRNAs) consist of closed loops of single-stranded RNA that are highly stable and abundant within the brain, particularly within synapses (Rybak-Wolf et al., 2015; You et al., 2015). As a result of their unique structure, circRNAs are resistant to exonuclease-mediated RNA degradation and are long-lived, exhibiting a median half-life at least 2.5 times greater than linear counterparts (Enuka et al., 2016; Jeck et al., 2013). They are also highly conserved and abundant within the brains of many organisms, are implicated in neurodegenerative and neuropsychiatric disorders, and exhibit dynamic regulation across many different cell-types, developmental stages, and neural activity (Jeck et al., 2013; Knupp & Miura, 2018; Lukiw, 2013; Rybak-Wolf et al., 2015; Wang et al., 2018; You et al., 2015; Zimmerman et al., 2020). Given all of the above features, particularly their stability, circRNAs are widely speculated to act as “memory molecules”. Yet, there are still relatively few studies directly linking circRNA function with memory.

One particularly robust memory protocol with relevance to human health is Pavlovian fear conditioning and fear extinction, which is behaviourally analogous to exposure therapy in humans and has well-established neuronal circuitry. For instance, the medial prefrontal cortex (mPFC) is a region that is critical for fear extinction memory and is comprised of the prelimbic prefrontal cortex (PLPFC) and the infralimbic prefrontal cortex (ILPFC), which mediate fear expression and fear extinction memory consolidation and retrieval, respectively (Sierra-Mercado et al., 2011; Stern et al., 2014; Thompson et al., 2010). Therefore, in order to establish the role of circRNAs in fear extinction memory, we decided to: 1) profile circRNAs in the mPFC of fear extinction-trained mice in a compartmentalised manner, and 2) manipulate the expression of a synapse-enriched circRNA within the neural processes of the ILPFC prior to fear extinction training.

Since circRNAs often originate from gene bodies and therefore share an overlapping sequence with the linear isoforms that are generated from the same locus, their identification is reliant on the detection of their back-splice junction (BSJ), which occurs at the site where a downstream 5’bss (back-splice site) and 3’ss (splice site) are covalently joined (Gao & Zhao, 2018; Jeck & Sharpless, 2014; Szabo & Salzman, 2016). A recent method called RPAD (RNAse R treatment, polyadenylation, and poly(A)+ depletion) was recently shown to improve circRNA detection by polyadenylating all exonucleolytically-digested RNAs with a 3’end (i.e., non-circular RNA) prior to performing poly(A)+/- selection to enrich for non-polyadenylated RNA (i.e., circular RNA) (Pandey et al., 2019a). As circRNAs are known to be enriched within synapses, a circRNA enrichment method that would be compatible with low-input material from both total cell (T) and synaptosome (S) compartments was required. Thus, it was decided to employ a modified version of the RPAD method, namely without RNAse R treatment as it requires relatively large RNA input, can be highly variable, and is known to deplete some circRNAs (Szabo & Salzman, 2016; Vincent & Deutscher, 2006; Y. Zhang et al., 2016).

In order to manipulate circRNA expression, the CRISPR-Cas-inspired RNA targeting system (CIRTS) was chosen due to its smaller size and mammalian origin versus the CRISPR-Cas13 targeting system of bacterial origin (Rauch et al., 2019). The CIRTS construct was then further modified so that it could specifically degrade circRNA targets within neural processes. Consequently, with these profiling and manipulation tools in hand, this study sought to establish a foundation for investigating circRNAs in the context of fear extinction memory.

## 2. Materials and methods

### 2.1. Animals

Animals (male C57BL/6J mice, 10-12 weeks old) were pair-housed using a plexiglass divider, maintained on a 12h light/dark cycle, and given free access to food and water. All behavioural testing was performed during the light phase in red-light conditions. All animal procedures were approved by The University of Queensland Animal Ethics Committee and conducted in accordance with the Australian Code of Practice for the Care and Use of Animals for Scientific Purposes.

### 2.2. cDNA synthesis and qPCR

Complementary DNA (cDNA) synthesis was performed using the QuantiTect Reverse Transcription Kit (Qiagen, #205313). Quantitative PCR (qPCR) was performed on a Qiagen Rotor-Gene Q 2plex real-time PCR cycler with 2x SensiFAST™ SYBR No-ROX mix (Bioline, #98020). Phosphoglycerate kinase 1 (*Pgk1*) and *18S rRNA* were used as internal controls (see Appendix A Table A.1 for primer pairs). All RNA levels for each PCR reaction were normalised relative to *Pgk1* or *18S rRNA* using the 2^-ΔΔCT^ method. For experiments using RNAse R treatment, the 2^-ΔCT^ method was used to normalise each treated sample with its corresponding Input (i.e., RNAse R-). Every PCR reaction was duplicated across two different runs.

### 2.3. CIRTS constructs

The following gRNA sequence was cloned into the BsmBI site of a modified CIRTS construct (**Fig. 2A**) in order to degrade *Cdr1as* (30nt reverse complementary to BSJ): GGCCAGATCTGAGCCTGGGAGCTCTCTGGCCTTATT**GCACCACTGGAAACCCTG GATACGGCAGAC**. A gRNA targeting red fluorescent protein (RFP), which is not present in neurons, was used as a genetic manipulation control (30nt reverse complementary to *mRuby3*): GGCCAGATCTGAGCCTGGGAGCTCTCTGGCCTTATT**GATACATCATCTCTGTAT TAGGCTCCCAAC**.

### 2.4. CircRNA manipulation and behavioural analysis

Adult male C57BL/6J animals (10-12 weeks old) were housed in split-cages for at least a week before implantation of double cannulae (PlasticsOne) targeted to the ILPFC (AP: +1.8mm, ML: 0mm, DV: -2.85mm). All animals were given at least a week to recover before behavioural training. Behavioural training took place in conditioning chambers (Coulborn Instruments) with two transparent walls, two stainless steel walls, and steel electric grid floors that were customised into two different contexts: Context A (CTX A) and Context B (CTX B). CTX A is used for fear conditioning (FC) where animals are placed directly onto the electric grid and the environment is scented with a 6% lemon extract/14% of 80% ethanol/water mix. CTX B is used for fear extinction training (EXT) where animals are placed inside a Perspex cylinder located on top of a transparent piece of square plastic that is covering the electric grid and the environment is scented with a 20% vinegar/20% of 80% ethanol/water mix. Cameras located above the set-up captured movement and automatically processed it using a freezing measurement program (FreezeFrame). The FC protocol consisted of 3x CS-US pairings of a 2-min, 80 dB, pure 16000Hz tone (CS) co-terminating with a 1s 0.7 mA foot shock (US) in CTX A. Animals were exposed to CTX A for 2 mins (pre-CS) prior to the first presentation of CS (each CS duration: 2 min) with an intertrial interval (ITI) of 2 mins between each CS presentation as well as 2 mins following the last CS presentation (post-CS) before animals were removed and placed back into their home cages. Animals were then assigned into their respective genetic manipulation (i.e., RFP or *Cdr1as*-targeting CIRTS constructs) and behavioural groups (i.e., RC or EXT) based on average freezing scores. The day after FC, transient genetic manipulation was achieved by infusing CIRTS constructs into the ILPFC using the *in vivo* jetPEI reagent (Polyplus, #101000040) for 2 days (2 µL/day) with an infusion rate of 0.2 µL/min administered using a 10 µL Hamilton glass syringe (Harvard Apparatus, #72-1774) attached to a PHD ULTRA™ syringe pump (Harvard Apparatus, #70-3005) (see **Fig. 2D** for behavioural timeline). After two days of infusions, animals were placed into CTX B and either subjected to repeated presentations of CS (e.g., 60CS) for EXT or left in CTX B for the same amount of time with no CS to act as a retention control (RC) that only has experience with the CS in an aversive setting. Pre-CS, Post-CS and CS durations were once again all 2 mins but the ITI between CS presentations was 5 seconds. The percentage of time the animal remains immobile (%Freezing) is used as a proxy for measuring the strength of the original fear memory and the subsequent RC/EXT training (FreezeFrame). The day after RC/EXT training, animals were placed back into CTX B and exposed to 3x CS presentations (same timing as FC but without US) to examine the strength of the fear extinction memory. The following day, animals were placed back into CTX A to investigate effects on fear renewal (3x CS). Six days later, animals were once again subjected to CTX B and CTX A tests over two days to investigate the strength of the extinction training and its consolidation a week later.

### 2.6. circRNA profiling

Samples for circRNA profiling were prepared by: 1) FC and 60CS RC/EXT training, 2) immediate mPFC extraction and pooling (4x mPFCs/biological replicate) following RC/EXT, 3) homogenisation, removal of total cell aliquot, and synaptosome isolation with the rest of the homogenate (see Section 2.8. for detailed methodology), and 5) circRNA enrichment. CircRNA enrichment is based on the RPAD method (Pandey et al., 2019b) but without RNAse R treatment. Briefly, total RNA was extracted according to NucleoZOL (Macherey-Nagel, #740404.200) manufacturer instructions and immediately processed as follows: 1) DNAse treatment to remove any trace of DNA (Invitrogen, #AM1907), 2) Poly(A)-tailing to add adenosines to the open 3’ ends of RNA (Lucigen, #PAP5104H), and 3) Poly(A) selection using Dynabeads™ Oligo(dT)_25_ beads (Invitrogen, #61005) to separate RNA into two populations, non-polyadenylated RNA (i.e., circular RNA) and polyadenylated RNA (i.e., ‘linear’ RNA with open 3’ ends). RNA input into circRNA enrichment was standardised to 300ng for both total cell and synaptosome samples. Following circRNA enrichment, libraries were prepared using the SMARTer® Stranded Total RNA-seq Kit v2 – Pico Input Mammalian (Takara, #634413). Samples were fragmented for 4 min @ 94°C and final amplification was performed using 16 cycles. After library preparation, libraries were size selected in order to have an average size of ∼400 bp using a 0.6x and 1.6x bead clean-up. Paired-end libraries were sequenced using the Illumina HiSeq 4000 sequencing platform (Genewiz) with read length of 150 bp*2. BWA-mem (v0.7.17) (Li & Durbin, 2009) was used for the alignment against mouse genome reference (mm10), and then CIRIquant (v1.1.1) (J. Zhang et al., 2020) was used to identify circular RNAs in each sample. For the differential expression analysis between groups, a customed analysis pipeline of edgeR was applied by utilizing the “prep_CIRIquant” script in the CIRIquant package for the generation of a matrix of circRNA expression level and the “CIRI_DE_replicate” script for the detection of differentially expressed circRNAs.

### 2.7. RNAse R treatment

RNAse R (Lucigen, #RNR07250) treatment was performed on 1 μg total RNA for 10 minutes at 37°C using 5U of enzyme. In order to improve RNAse R processivity through structured regions of RNA (e.g., G-quadruplexes, rRNA), a LiCl-containing buffer (0.2M Tris-HCl (pH 8), 1M LiCl, 1mM MgCl_2_) was used instead of the original KCl-containing buffer (0.2M Tris-HCl (pH 8), 1M KCl, 1mM MgCl_2_) supplied with the enzyme (Xiao & Wilusz, 2019).

### 2.8. Synaptosome isolation

Synaptosomes were isolated from fresh tissue immediately following behavioural training using a discontinuous Percoll-sucrose density gradient with modifications. For each synaptosome isolation, 4x mPFCs were homogenised in 2 mL of ice-cold Gradient Medium (GM) buffer (250mM sucrose, 5mM Tris-HCl (pH 7.5), Milli-Q) with freshly added RNAseOUT (Invitrogen, #10777019, 1:1000) and DTT (5mM final concentration). 100 µL of homogenate was set aside as total cell (T) and mixed with NucleoZOL (Macherey-Nagel, #740404.200) before storage at -80°C. The rest of the homogenate (∼1900 µL) was spun at 1000 x g for 10 min at 4°C to pellet cellular debris and nuclei. The supernatant was then slowly added to a discontinuous Percoll gradient consisting of 3%, 10%, and 23% Percoll (GE Healthcare, #17-0891-01) layers (6 mL/layer) in GM buffer. For extra material, the pellet was resuspended in 2 mL of GM buffer and centrifuged again. The samples were then spun at 31000 x g (18468 rpm) for 5 min at 4°C in a fixed-angle rotor (50.2 Ti rotor in Beckman Coulter Optima L-100 XP) with slow brake. Following centrifugation, layers F1/2 were removed before transferring the combined F3/4 layer into a clean 25mL centrifuge tube (Beckman Coulter, #355618) and diluting in GM buffer (at least 1:4) before centrifuging again at 20000 x g (or 14834 rpm, 50.2 Ti rotor) for 30 min at 4 °C, with slow brake. Afterwards, supernatant was removed carefully and the synaptosome pellet (∼1mL) was transferred to a 2mL tube and topped up with GM buffer before centrifugation at 18000 x g for 10 min at 4°C. Finally, after removing as much supernatant as possible, the pellet was resuspended in 500 µL of NucleoZOL (Macherey-Nagel, #740404.200) before storage at - 80°C.

### 2.9. Immunostaining

Adult male C57BL/6J mice were intracardially perfused using ice-cold PBS to wash out the blood before switching to ice-cold fixative 4% paraformaldehyde in PBS. Brains were extracted and left to fix overnight in 4% paraformaldehyde in PBS at 4°C before washing the brains a few times with PBS for a couple mins/wash. Brains were then sectioned into 40-50µm slices using a vibratome and either stored at 4°C in PBS (short-term storage) or 0.01% NaN_3_ in PBS (long-term storage). For visualising the correct infusion and expression of CIRTS constructs (**Fig. 2B**), slices were first blocked using blocking solution (5% normal goat serum (Sigma, #G9023, Lot:SLCH5798) in 0.3% Triton X-100 in PBS) for 1 hour at room temperature prior to overnight incubation on a shaker in primary antibody solution (1:2000 Rabbit α GFP diluted in blocking solution, Abcam, #ab6556, Lot: GR3351352-1, preincubated for 15 minutes prior to adding to sections) at room temperature. Afterwards, sections were washed 3x in 0.3% Triton X-100 for 15 mins/wash before incubating in secondary antibody solution (1:1000 Goat α Rabbit, Invitrogen, #A11034, Lot: 1812166 in blocking solution) for 1 hour. Afterwards, sections were washed 3x in 0.3% Triton X-100 in PBS for 15 mins/wash before incubating in DAPI solution (1:2000 DAPI, Invitrogen, #D1306, Lot:1942279 in 0.3% Triton X-100 in PBS) for 10 mins at room temperature and then washed 1x in PBS for 5 minutes before mounting onto 26×76mm glass microscope slides (Thermo Scientific™ SuperFrost™, #MIC3020) using fluorescence mounting media (Dako, #S3023) and 24×60mm coverslips (Menzel, #1.5 thickness). Rabbit IgG (1:4000 Cell Signaling Technology, #2729S, Ref: 11/2016) was used as a control to visualise background staining caused by non-specific IgG binding. Images were acquired using an Axio Imager Z1 upright fluorescence microscope fitted with an Axiocam MRm camera (Carl Zeiss Pty Ltd) using a 10x objective with numerical aperture 0.3 (0.52mm WD, 0.645 μm/pixel) and ZEN software (Carl Zeiss). All images were converted into RGB tiffs using ImageJ and then further processed in Adobe Photoshop to threshold for background fluorescence staining using the IgG control sections as a baseline.

### 2.10. Statistical analyses

For qPCR experiments (**Fig. 1E-F**), unpaired t-tests with Welch’s correction were performed relative to their respective control group. For behavioural experiments (**Fig. 2E-F**), a two-way ANOVA was performed with Dunnett’s post hoc tests relative to RFP RC control animals. All alpha levels were set to 0.05.

**Figure 1:**
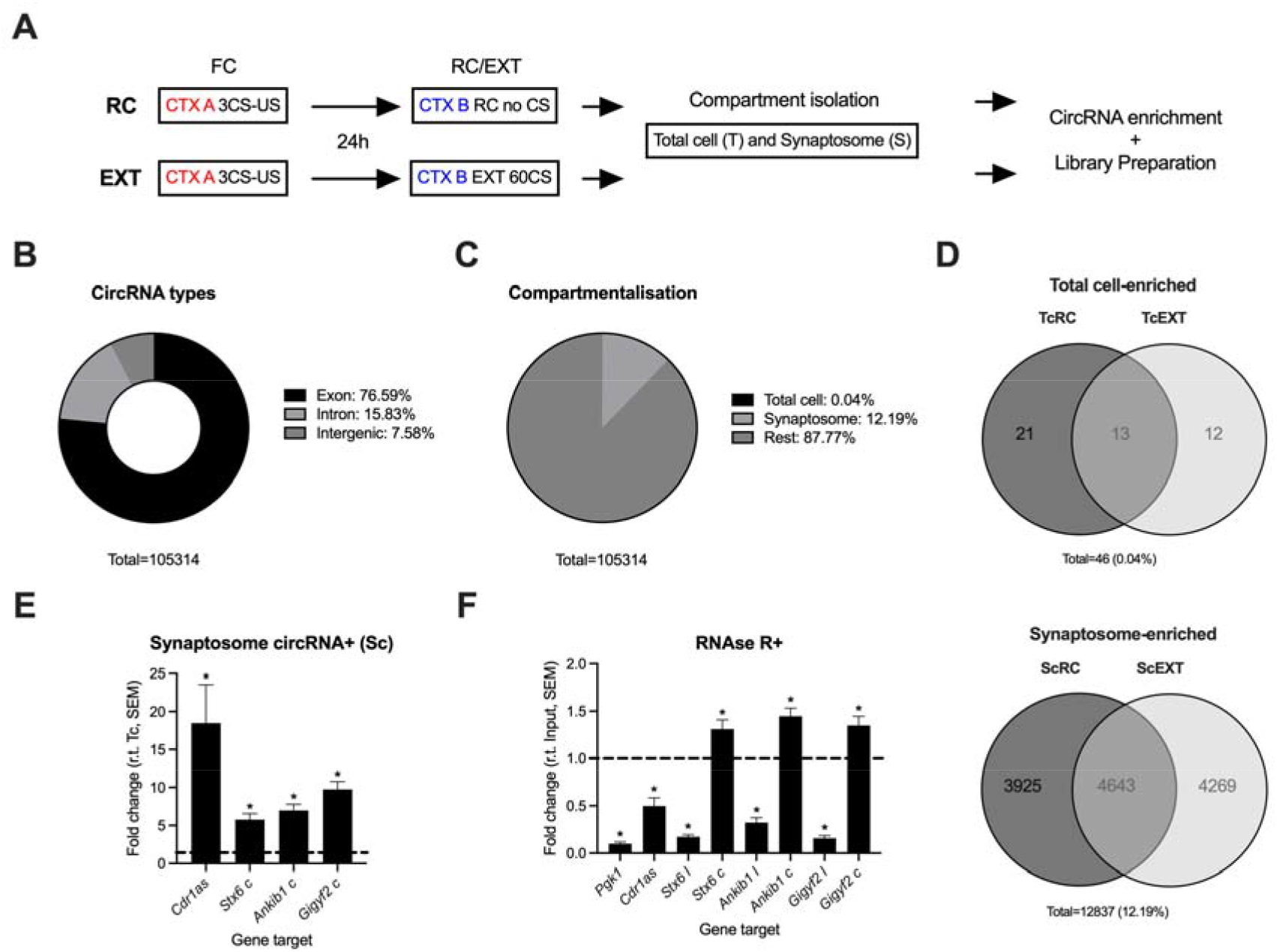
Compartmentalised profiling (total cell vs synaptosome) revealed thousands of circRNAs enriched in the neural processes of the mPFC of adult male C57BL/6J mice after RC/EXT training. **A)** Timeline of circRNA profiling: behaviour, compartment isolation, circRNA enrichment, and library preparation. Animals were first fear conditioned and then assigned into biological replicates (*n* = 4 animals/biological replicate) based on average freezing score prior to 60CS RC/EXT training and fresh synaptosome isolation followed by circRNA enrichment and library preparation. Illumina short-read sequencing followed by BSJ detection was used to identify circRNA candidates. **B)** Most circRNAs identified in this study were exonic in origin followed by intronic and intergenic. **C)** The majority of circRNAs identified were not significantly enriched in either the total cell or synaptosome fraction (87.77%, fold change < ± 1 and FDR > 0.05). A relatively higher proportion of circRNAs were enriched in the synaptosome (12.19%) compared to those enriched in the total cell compartment (0.04%). Synaptosome-enriched: 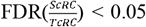 and 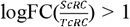 **AND/OR** 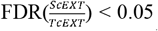 and 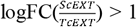. Total cell-enriched: 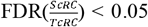 and 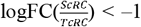 **AND/OR** 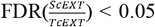 and 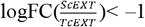. **D)** Breakdown of the total cell- and synaptosome-enriched fractions into behavioural groups RC and EXT. Overlap is the set of circRNAs that are enriched in their respective compartment in both RC and EXT groups. **E)** Graph depicting the fold change of target circRNAs (*Cdr1as, Stx6 c, Ankib1 c*, and *Gigyf2 c*) in the synaptosome circRNA+ (Sc) population relative to the total cell circRNA+ (Tc) population. All values are calculated using the 2^-ΔΔCT^ analysis method using *Pgk1* as a housekeeping gene with each Sc sample normalised to its respective Tc sample. The dashed line at 1 denotes the Tc threshold line where values above the line are increased relative to Tc (i.e., synaptosome enriched) and those below are decreased relative to Tc (i.e., total cell enriched). *n* = 12/column. **F)** RNAse R treatment was used to validate the circularity of circRNAs that originate from the genes *Stx6, Ankib1*, and *Gigyf2* alongside their linear counterparts, denoted ‘c’ and ‘l’ respectively. All values are calculated using the 2^-ΔCT^ analysis method with each sample normalised to its respective Input (RNAse R-samples), which is denoted by the dashed line at fold change = 1. *Pgk1* is a positive linear RNA control to demonstrate that the RNAse R enzyme is functional. If an RNA is linear, then it will be decreased relative to Input (i.e., below the dashed line); if an RNA is circular, it will be increased relative to Input (i.e., above the dashed line). *n* = 9/column. *Cdr1as* is a well-known circRNA, however, it is known to be sensitive to RNAse R treatment (Cheng et al., 2016; Szabo & Salzman, 2016). Asterisks on the graphs (E-F) indicate significance level using unpaired t-tests with Welch’s correction relative to Tc (E) and Input (F): *P<0.05. All error bars depict SEM. *Abbreviations: Ankib1*, Ankyrin Repeat and IBR Domain Containing 1; BSJ, back-splice junction; c = circRNA, circular RNA; *CDR1*, Cerebellar Degeneration Related Protein 1; *Cdr1as*, CDR1 antisense RNA; CS, conditioned stimulus; CTX, context; EXT, extinction; FC, fear conditioning *or* fold change if in “logFC”; FDR, false discovery rate; *Gigyf2*, GRB10 Interaction GYF Protein 2; l = linear RNA; mPFC, medial prefrontal cortex; RC, retention control; RNA, ribonucleic acid; RNAse, ribonuclease; r.t., relative to; S, synaptosome; Sc, synaptosome circRNA+ population; SEM, standard error of the mean; *Stx6*, syntaxin 6; T, total cell; Tc, total cell circRNA+ population; US, unconditioned stimulus.

## 3. Results and discussion

### 3.1 Profiling circRNAs associated with fear extinction within the mPFC of adult male C57BL/6J mice in a compartmentalised manner

For this experiment, adult male C57BL/6J mice (10-12 weeks old) were fear conditioned and then divided into either the RC or EXT group (**Fig. 1A**). A full 60CS EXT paradigm, alongside the RC group with no CS presentations, was performed and the mPFCs of trained mice were extracted immediately following behaviour and processed for synaptosome isolation (*n* = 4x mPFCs/biological replicate, 6 biological replicates/group). Following this, samples underwent circRNA enrichment and library preparation before sequencing on the Illumina platform (see Section 2.6.). Bioinformatic analysis revealed that the majority of circRNAs detected were exonic in origin followed by intronic and intergenic (**Fig. 1B**). A small proportion (0.04%) were significantly enriched in the total cell fraction versus synaptosome whilst a slightly higher proportion of circRNAs were significantly enriched in the synaptosome fraction (12.19%) compared to total cell (**Fig. 1C-D**, see Appendix B).

Using an independent biological cohort, the findings of the sequencing analysis were replicated using divergent PCR (D-PCR) to show that circRNA targets from *Ankib1, Gigyf2*, and *Stx6* (denoted *Ankib1 c, Gigyf2 c*, and *Stx6 c* respectively, see Appendix A Table A.2) as well as the well-known circRNA *Cdr1as* were indeed enriched in the synaptosome fraction (**Fig. 1E**). Due to the removal of RNAse R treatment from the circRNA enrichment approach, RNAse R treatment was used in combination with D-PCR as an orthogonal method to validate the circularity of these targets (**Fig. 1F**). *Pgk1* was included as a known linear RNA (negative control) whilst *Cdr1as* is a well-known circRNA (positive control). However, despite its known circularity, the data shows that *Cdr1as* is sensitive to RNAse R treatment and is decreased relative to Input, similar to what is expected from a linear RNA. This false negative result is not entirely unexpected since other studies have also noted that RNAse R is able to degrade some circRNAs, including *Cdr1as*, which may be due to its relatively large size as larger circRNAs are more likely to be nicked and linearised compared to smaller circRNAs (Cheng et al., 2016; Jeck et al., 2013; Rahimi et al., 2021; Szabo & Salzman, 2016; Y. Zhang et al., 2016).

Interestingly, there were only a handful of circRNAs with statistically significant differences in abundance between RC and EXT (24); however, after eliminating candidates with low abundance and/or more than a couple of replicates with no BSJ detection, there were none (0) (Appendix B). Across the entire dataset, each circRNA candidate exhibited large variability between replicates in the number of BSJs detected, which could have obscured any real differences between RC and EXT behavioural groups. The observed variability could be reflective of the natural biological variance of circRNAs between animals or perhaps the variance between replicates was introduced artificially during pooling (4x mPFCs = 1x replicate) and/or circRNA enrichment and subsequent processing (e.g., enzymatic reaction variability, library preparation, etc.). Since bulk mPFCs were pooled from four animals in order to perform synaptosome isolation, this could mean that some replicates may have contained animals that did not learn as well as others.

An improved approach would be to isolate circRNAs from the tagged engram cells of an individual animal. However, this would mean sacrificing compartment resolution (i.e., synaptosome isolation) and there also may not be enough total RNA yield to perform circRNA enrichment. This is because a lot of circRNAs are present at the synapse and fluorescence-activated cell sorting (FACS) sorting shears off these synaptic compartments, which makes FACS better suited to studies of nucleic acid in the nucleus. Moreover, it is important to note that RNA abundance does not necessarily correlate with its function; perhaps the circRNAs are being functionally modified or recruited during extinction learning and are interacting with other molecules within the cell to exert their effects. Additionally, the number of BSJs detected for a given circRNA is not truly reflective of its ‘abundance’ due to PCR amplification during library preparation and the limitation that circRNAs can only be identified by their BSJ. In any case, if it is accepted that there are no significant differences in the abundance of circRNAs between RC and EXT groups, the proposal that circRNAs are immediately expelled from the cell during fear extinction learning as a messenger between cells or to rapidly alter their levels at the synapse is weakened. Multiple approaches and improved cellular resolution will likely be required to uncover the true functional differences of circRNAs in fear extinction memory.

### 3.2. Knockdown of Cdr1as in the ILPFC of adult male C57BL/6J mice impairs fear extinction memory

*Cdr1as* has previously been shown to dampen neural activity with *Cdr1as* KO mice exhibiting increased spontaneous vesicle release as well as stronger synaptic depression following enhanced neural activity in excitatory neurons, all of which is thought to contribute to its sensorimotor gating deficit, a phenotype that is associated with neuropsychiatric disorders such as schizophrenia (Piwecka et al., 2017). Yet, despite its effect on neural activity, *Cdr1as* knockout (KO) mice showed no effect on recognition memory (Piwecka et al., 2017). Since it was a KO mouse model, there is the possibility that other mechanisms were recruited to compensate for the loss of *Cdr1as*. Therefore, we decided to transiently knockdown (KD) *Cdr1as* expression in the adult mouse brain to see if this would have any effect on fear extinction learning and memory.

Since *Cdr1as* is abundant within the neural processes, a neuron-specific (synapsin I-driven) CIRTS construct with a Pin nuclease for degradation that is directed towards the synapse for local translation via a *Calm3* intron localisation signal (Sharangdhar et al., 2017) was designed to knockdown the expression of *Cdr1as* in the neural processes of the ILPFC by targeting the 30nt centred around its BSJ using antisense gRNAs expressed off the same construct (**Fig. 2A-C**) (Rauch et al., 2019). RFP, which is not naturally present in wild-type C57BL/6J mice, was used as the control target. The RFP-and *Cdr1as*-targeting CIRTS constructs were transiently transfected into the ILPFC of adult male C57BL/6J mice following fear conditioning using the *in vivo* transfection reagent jetPEI (Polyplus, #101000040) (**Fig. 2D**). This approach led to the impairment of fear extinction memory in animals treated with the *Cdr1as*-targeting CIRTS construct, which also generalised to CTX A (**Fig. 2E-F**, see Appendix A Fig A.1 for full behavioural data). Moreover, since transfection only lasts for a few days using the transfection reagent jetPEI (Polyplus, #101000040), the *Cdr1as*-targeting CIRTS construct appears to have affected the actual formation/consolidation of fear extinction memory and not just its expression during recall.

**Figure 2:**
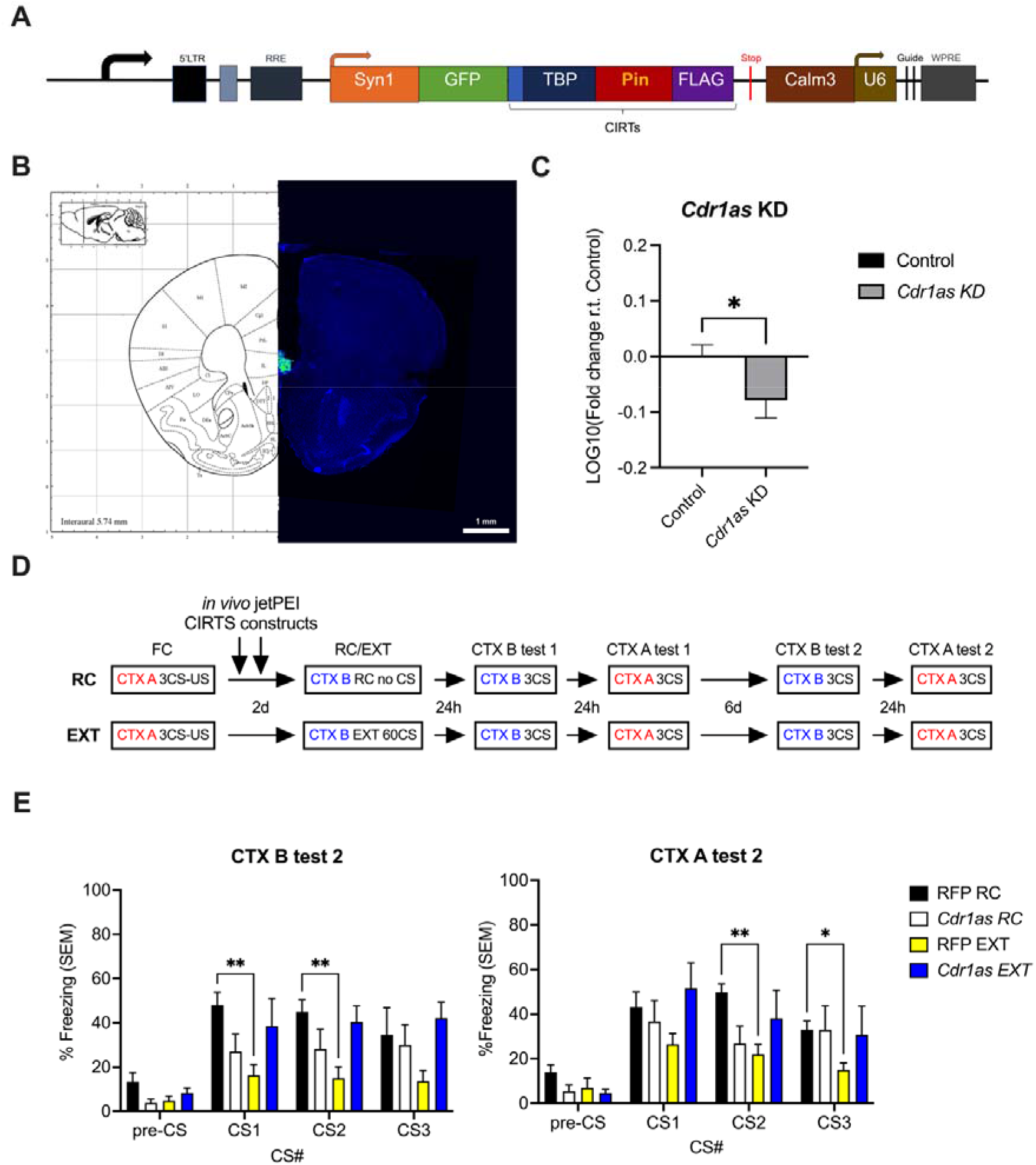
Knockdown of *Cdr1as* in the neural processes of the ILPFC impairs fear extinction memory. **A) T**he CRISPR-Cas-inspired RNA targeting system (CIRTS) (Rauch et al., 2019) is comprised of a gRNA and several protein parts. The gRNA is composed of a sequence that is complementary to the target of interest and a TAR hairpin. The RNA hairpin binding protein (TBP), which binds the TAR hairpin, a ssRNA binding protein, and an effector domain are linked together. The effector domain can be swapped out depending on the desired effect (e.g., degradation, translation, or editing). The modifed CIRTS plasmid illustrated here was generated by cloning the CIRTS protein parts and guides into a pFSy(1.1)GW plasmid backbone (Addgene, #27232) (Dittgen et al., 2004) with a synapsin I promoter (neuron-specific). A GFP protein was fused to the modular CIRTS protein domains via a 2A self-cleaving peptide signal (T2A) for visualisation and a *Calm3* intron localisation signal (Calm3_L_3’UTR) (Sharangdhar et al., 2017) was appended to the end to promote localisation to the neural processes upon neural activation. The Pin nuclease is the effector domain used to degrade targets complementary to the gRNA, which is driven off the same construct under the U6 promoter by cloning into the BsmBI site. **B)** Image depicting a representative example of correct cannula placement and targeting of the CIRTS constructs (green) to the ILPFC of an adult male C57BL/6J mouse. Scale bar: 1 mm. **C)** Infusion of the modified CIRTS construct targeting the *Cdr1as* BSJ led to a small but significant knock-down in *Cdr1as* expression. Asterisk on the graph indicates significance level using an unpaired t-test with Welch’s correction relative to Control (animals that were either uninfused or infused with RFP-targeted CIRTS construct at both 60CS EXT and baseline): *P<0.05. *n* = 17-19/group. All error bars depict SEM. **D)** Behavioural timeline for the experiment investigating the effect of circRNA *Cdr1as* in fear extinction memory. *Cdr1as* was transiently knocked down in neurons of the ILPFC after fear conditioning and prior to extinction training using *in vivo* jetPEI transfection of the *Cdr1as*-targeting CIRTS construct shown in A), with an RFP-targeting CIRTS construct used as a control. **E)** The strength of the extinction memory was investigated by placing animals back into CTX B and CTX A in the days immediately following RC/EXT training (CTX B/A tests 1, not shown) and a week later (CTX B/A tests 2). Context recall tests were performed using 3x CS presentations (see Section 2.4. for more details). The amount of time they spent freezing in each context (%Freezing) is used to measure how well they recall their training; a high freezing score indicates more fear whereas a low freezing score indicates less fear. For EXT animals, it is expected that they will have less freezing compared to RC controls. RFP EXT shows significant differences to RFP RC controls whereas *Cdr1as* EXT shows impaired recall of fear extinction memory. For E)-F), the two-way ANOVA results for group (outlined in figure legend) is as follows: E) F_3,18_=3.919, *P=0.0258, F) F_3,18_=1.736, P=0.1954. Asterisks on the graphs indicate significance level using Dunnett’s post hoc tests relative to RFP RC control animals: *P<0.05, **P<0.01. *n* = 5-7/group. All error bars depict SEM. *Abbreviations:* 5’LTR, 5’ long terminal repeat, which acts as a RNA Pol II promoter; BSJ, back-splice junction; Calm3, Calm3_L_ 3’UTR sequence, which localises RNA to dendrites via Staufen2; *CDR1*, Cerebellar Degeneration Related Protein 1; *Cdr1as*, CDR1 antisense RNA; CIRTS, CRISPR-Cas-inspired RNA targeting system; CS, conditioned stimulus; CTX, context; d, days; EXT, extinction; FC, fear conditioning; FLAG, DYKDDDDK-tag; GFP, green fluorescent protein; h, hours; KD, knockdown; Pin, Pin nuclease; RFP, red fluorescent protein; RC, retention control; RRE, Rev Response Element; r.t., relative to; SEM, standard error of the mean; Syn1, synapsin I promoter; TBP, TAR-binding protein hairpin binding domain; U6, U6 promoter for driving small hairpin RNA expression; US, unconditioned stimulus; WPRE, woodchuck hepatitis virus post-transcriptional regulatory element.

### 3.3. Changes in the Cdr1as ncRNA network

*Cdr1as* has previously been shown to be a member of a ncRNA network with *miR-7, miR-671*, and the lncRNA *Cyrano* (Kleaveland et al., 2018). *Cdr1as* contains >70 partial *miR-7* binding sites (130 in mice, 73 in human) with one almost perfect complementary binding site for *miR-671* (Hansen et al., 2013; Kleaveland et al., 2018; Piwecka et al., 2017). These features enable *Cdr1as* to be targeted by *miR-671* for degradation as well as to protect *miR-7* from degradation by *Cyrano*. Unchecked *miR-7* enhances *miR-671*-directed cleavage of *Cdr1as* whilst the lncRNA *Cyrano* regulates *Cdr1as* accumulation by degrading *miR-7* (Kleaveland et al., 2018). In mice, three *miR-7* loci express two different variants: *miR-7a* (2) and *miR-7b* (1). *miR-7* targets include transcripts involved in synaptic plasticity, cytoskeletal organisation, and vesicle fusion and trafficking (Hayashi et al., 2004; Latreille et al., 2014; Memczak et al., 2013; Zhao et al., 2020).

In order to investigate changes in the *Cdr1as* ncRNA network, the expression levels of *Cdr1as, Cyrano*, and *miR-7* precursors (pri-miR-7a1, pri-miR-7a2, pri-miR-7b) were profiled during the course of RC/EXT training (10CS, 30Cs, 60CS) as well as after *Cdr1as* KD (**Fig. 3**). *Cdr1as* levels decreased from the beginning of training (10CS) to the end (60CS), which could be indicative of increased degradation by *miR-671* and the subsequent release of *miR-7* into the cellular milieu. During RC/EXT training, the *miR-7* precursor pri-miR-7a2 is upregulated at 30CS compared to RC10 before returning to baseline by 60CS (**Figure 13A**). *Cyrano* and *miR-7* precursors, pri-miR-7a1 and pri-miR-7b, show no significant changes in expression levels compared to RC10. However, during *Cdr1as* KD conditions, there is an upregulation in *miR-7* precursor pri-miR-7a1 with no significant changes in the other *miR-7* precursors and *Cyrano*.

**Figure 3:**
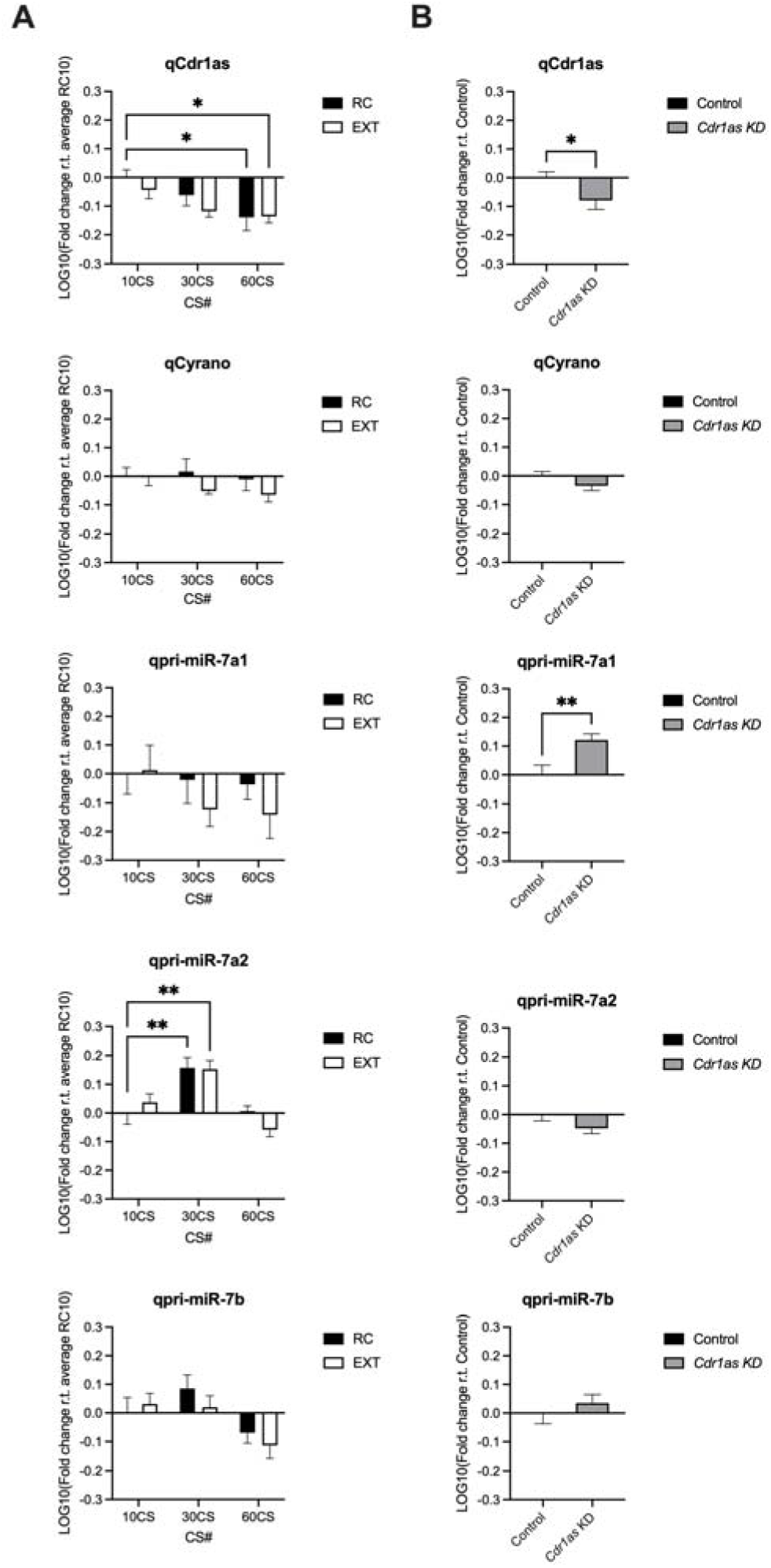
Changes in the *Cdr1as* ncRNA network under learning conditions and following *Cdr1as* KD. **A)** The expression levels of circRNA *Cdr1as*, lncRNA *Cyrano*, and *miR-7* precursors (pri-miR-7a1, pri-miR-7a2, and pri-miR-7b) within the mPFC of adult male C57BL/6J mice sacrificed at 10CS, 30CS, and 60CS timepoints of RC/EXT training are shown relative to RC10. All values were calculated using the 2^-ΔΔCT^ analysis method, using *Pgk1* as the housekeeping gene and the average RC10 ΔCt as the normalisation value, before transformation using LOG10. A two-way ANOVA with multiple comparisons made using a post-hoc Dunnett’s test with RC10 as the control cell for comparison was used. *P<0.05, **P<0.01. *n* = 8/group. All error bars depict SEM. **B)** The effect of *Cdr1as* KD within the ILPFC of adult male C57BL/6J mice on the expression levels of circRNA *Cdr1as*, lncRNA *Cyrano*, and *miR-7* precursors (pri-miR-7a1, pri-miR-7a2, and pri-miR-7b) was investigated by performing qPCRs using cDNA samples obtained from animals sacrificed 1 day following *Cdr1as* KD treatment (*in vivo* jetPEI transfection) at both baseline and immediately following EXT60 training. All values were calculated using the 2^-ΔΔCT^ analysis method, using *Pgk1* as the housekeeping gene and the average Control ΔCt as the normalisation value, before transformation using LOG10. An unpaired t-test with Welch’s correction was used to compare *Cdr1as* KD samples to the Control group (uninfused or infused with RFP-targeted CIRTS construct). *P<0.05, **P<0.01. *n* = 15-19/group. All error bars depict SEM. All primer sequences, except qCdr1as, were sourced from (Kleaveland et al., 2018) (A, B). *Abbreviations: CDR1*, Cerebellar Degeneration Related Protein 1; *Cdr1as*, CDR1 antisense RNA; circRNA, circular RNA; CS, conditioned stimulus; EXT, extinction; EXT60, 60CS EXT protocol; KD, knockdown; miR-X, miRNA X; miRNA, micro RNA; mPFC, medial prefrontal cortex; *Pgk1*, phosphoglycerate kinase 1; qX, qPCR primer for target X; qPCR, quantitative polymerase chain reaction; RC, retention control; RC10, retention control at 10CS; RFP, red fluorescent protein; RNA, ribonucleic acid; r.t., relative to.

The upregulation in pri-miR-7a2 expression during RC/EXT training may occur in order to: 1) replenish depleted *miR-7* levels so that it can be protected by *Cdr1as* once its expression levels recover in order to be ready for the next learning event, or 2) depending on how fast pri-miR-7a2 can be processed into mature *miR-7a*, trigger further degradation of *Cdr1as* and increases in the local concentration of *miR-7*. However, when *Cdr1as* is knocked down the day before, there is less *Cdr1as* available to protect *miR-7* and pri-miR-7a1 may be the primary loci responsible for maintaining *miR-7* levels. Thus, perhaps the pri-miR-7a2 locus is responsive to conditions of learning such as in the instance where *miR-7* release is triggered by *miR-671* degradation of *Cdr1as*, however, when *Cdr1as* levels are already lowered, the cell activates the pri-miR-7a1 locus to compensate for the increased turnover of *miR-7* due to less protection from *Cdr1as*.

In terms of the effect on *miR-7* targets during *Cdr1as* KD, it has previously been noted that there is a difference in the overall response of *miR-7* targets depending on if *Cdr1as* is removed conditionally (KD) versus constitutively (KO) (Piwecka et al., 2017). Following knockdown of *Cdr1as* in HEK293 cells there is increased repression of miR-7 targets, which is consistent with the idea that *Cdr1as* degradation leads to the sudden release of *miR-7*:RISC complexes within the cell (Memczak et al., 2013). In contrast, when *Cdr1as* is knocked out, there is an upregulation of *miR-7* targets and this is likely due to increased *miR-7* turnover due to a loss of *Cdr1as* protection (Piwecka et al., 2017). However, given the small effect sizes of up- and down-regulation of *miR-7* targets observed after largescale manipulation of *Cdr1as* and *miR-7*, those in the field have come to the hypothesis that *Cdr1as* acts as a vehicle to control the spatiotemporal delivery of *miR-7* and thereby controls the repression of *miR-7* targets involved in neural activity in a local manner rather than causing widespread repression (Kleaveland et al., 2018; Piwecka et al., 2017). Hence, *Cdr1as* acts as a regulatory brake on neural activity and the where and when of its activity and release of *miR-7* (i.e., local spine delivery) appears to be critical for its role in finetuning certain memory networks over others. Thus, when *Cdr1as* was knocked down in the neural processes the day prior to RC/EXT training in this study, *miR-7* local delivery chains were likely disrupted and this loss of precision in finetuning the balance of neural activity resulted in an impairment of fear extinction memory.

## 4. Concluding remarks

In this study, circRNAs within the mPFC of the adult mouse brain were profiled after fear extinction learning in both the total cell and synaptosome compartment using a modified version of the RPAD method. In doing this, 12837 circRNAs were found to be enriched at the synapse, including the well-known circRNA *Cdr1as*. Targeted knockdown of *Cdr1as* within the neural processes of the ILPFC of adult male C57BL/6J mice led to impaired fear extinction memory, which is likely the result of disrupted local delivery of *miR-7*. Altogether, these findings illustrate the importance of dynamic and localised circRNA activity at the synapse for memory formation, and further emphasises the importance of employing techniques that manipulate circRNA expression/function in a spatiotemporally controlled manner.

## Supporting information

Appendix B. Sequencing data

## CRediT authorship contribution statement

**Esmi Lau Zajaczkowski:** Conceptualization, Formal analysis, Investigation, Methodology, Project administration, Validation, Writing – original draft, Writing – review & editing. **Qiongyi Zhao:** Data curation, Formal analysis, Investigation, Software. **Wei-Siang Liau:** Investigation, Methodology, Validation. **Hao Gong:** Investigation, Methodology, Validation. **Sachithrani Umanda Madugalle:** Investigation, Methodology, Validation. **Ambika Periyakarrupiah:** Investigation, Visualization. **Laura Jane Leighton:** Investigation. **Mason Musgrove:** Investigation. **Haobin Ren:** Investigation. **Joshua Davies:** Investigation. **Paul Robert Marshall:** Conceptualization, Formal analysis, Investigation, Methodology, Supervision, Writing – review & editing. **Timothy William Bredy:** Conceptualization, Funding acquisition, Resources, Supervision, Writing – review & editing.

## Declaration of Competing Interests

The authors declare that they have no known competing financial interests or personal relationships that could have appeared to influence the work reported in this paper.

## Acknowledgements

The authors would like to thank Ms. Rowan Tweedale for helpful editing of the transcript.

## Funding

This research was supported by funding from the ARC (DP180102998-TWB), an Australian Government Research Training Program Scholarship and a Westpac Future Leaders Scholarship.

## Appendix A Supplementary material

The following are the supplementary material for this article: <u>Appendix A. Supplementary material.docx</u>

## Appendix B Sequencing data

The following are the sequencing data for this article: <u>Appendix B. Sequencing data.xlsx</u>

## Notes

### Competing Interest Statement

The authors have declared no competing interest.

