## Appendix A. Supplementary material for "Inhibiting circRNA *Cdr1as* expression in the ILPFC of adult male C57BL/6J mice impairs fear extinction memory"

**Table A.1:** Primer names and their corresponding sequences used throughout this study. D-PCR primer sequences were used for circRNA targets listed below, which only amplify the sequence surrounding the circRNA BSJ and result in no product for the linear RNA from the same gene. CircRNA BSJ coordinates (mm9) are shown below their designation. *Cdr1as is not located within a gene but is antisense to the CDR1 gene. ^#^Primer sequences for Cyrano and miR-7 precursors were obtained from: (Kleaveland et al., 2018). Abbreviations: Ankib1, Ankyrin Repeat and IBR Domain Containing 1; c = circRNA, circular RNA; CDR1, Cerebellar Degeneration Related Protein 1; Cdr1as, CDR1 antisense RNA; D-PCR, divergent PCR; F = forward primer; Gigyf2, GRB10 Interaction GYF Protein 2; l = linear RNA; lncRNA, long noncoding RNA; miR-7, miRNA 7; PCR, polymerase chain reaction; Pgk1, phosphoglycerate kinase 1; qX = qPCR primer against X; R = reverse primer; RNA, ribonucleic acid; rRNA, ribosomal RNA; Stx6, syntaxin 6.

| **Gene** | **Designation** | **Primer name** | **Sequence (5’ 🡪 3’)** |
| --- | --- | --- | --- |
| *Pgk1* | Linear | qPGK F | TGCACGCTTCAAAAGCGCACG |
|  |  | qPGK R | AAGTCCACCCTCATCACGACCC |
| *18S rRNA* | Linear | q18SrRNA F | CTGGATACCGCAGCTAGGAA |
|  |  | q18SrRNA R | GAATTTCACCTCTAGCGGCG |
| *Ankib1* | Circular  (chr5:3747021\|3772787) | qAnkib1 c F | CCCAGGATCTTCGTAGGTTAAAA |
|  |  | qAnkib1 c R | TTTGTTATTTTCTGATAGGCAGTGG |
|  | Linear | qAnkib1 l F | GGCGCATCCTCAAGTGTTCT |
|  |  | qAnkib1 l R | CCCTGATGATCTTGTGCCGA |
| *Gigyf2* | Circular  (chr1:89260679\|89276583) | qGigyf2 c F | TGACAGGCGCTTTGAAAAACCAG |
|  |  | qGigyf2 c R | AATTTATACTTTGGCAGGGCCGGA |
|  | Linear | qGigyf2 l F | TGATGCAGTGAAAGAGGTGGG |
|  |  | qGigyf2 l R | ACTGGGTAAATCCATCTTGGGC |
| *Stx6* | Circular  (chr1:157021116\|157044565) | qStx6 c F IV | GTCTCGACTGGACAACGTGA |
|  |  | qStx6 c R IV | TCTCTGAAACAATCCTTGGGCA |
|  | Linear | qStx6 l F | CTCGACTGGACAACGTGATGAA |
|  |  | qStx6 l R | TAGGAAGAGGATCAGCACGAC |
| *Cdr1as** | Circular  (chrX:58436423\|58439349) | qCdr1as F | CCAGTGTATCGGCGTTTTGAC |
|  |  | qCdr1as R | TCACGATTGTCTGGAAGACCTTGAC |
| *Cyrano^#^* | Linear  (lncRNA) | qCyrano F (mm_Cyrano_qPCR_F1) | ACCATGGCTCACCTTTATGC |
|  |  | qCyrano R  (mm_Cyrano_qPCR_R1) | TGGGGAAAGTAACAGGATGG |
| *miR-7^#^* | *mus musculus* miR-7 precursor: **pri-miR-7a1** | qpri-miR-7a1 F  (mm_pri-miR-7a1_qPCR_F1) | GCCTGTAGAAAATGTAGAAGAGAC |
|  |  | qpri-miR-7a1 R  (mm_pri-miR-7a1_qPCR_R1) | TATGGCAGACTGTGATTTGTTG |
|  | *mus musculus* miR-7 precursor: **pri-miR-7a2** | qpri-miR-7a2 F  (mm_pri-miR-7a2_qPCR_F1) | AAGTCAGGGGAGCAGGCAC |
|  |  | qpri-miR-7a2 R  (mm_pri-miR-7a2_qPCR_R1) | TGGCAGACTGGGACTTGTTGT |
|  | *mus musculus* miR-7 precursor: **pri-miR-7b** | qpri-miR-7b F  (mm_pri-miR-7b_qPCR_F1) | GAGAGAGAGAGAAGCACTTGAGGG |
|  |  | qpri-miR-7b R  (mm_pri-miR-7b_qPCR_R1) | GAGGCTGGCTGTGACTTGTTGT |

**Table A.2:** CircRNA targets chosen for further validation. The circRNA coordinates indicate the position of the BSJ with a ‘|’. Cdr1as is a well-known circRNA and serves as a positive control. circBase was used to find the estimated spliced length (Glažar et al., 2014). Abbreviations: Ankib1, Ankyrin Repeat and IBR Domain Containing 1; BSJ, back-splice junction; CDR1, Cerebellar Degeneration Related Protein 1; Cdr1as, CDR1 antisense RNA; c = circRNA, circular RNA; Gigyf2, GRB10 Interaction GYF Protein 2; ID, identification; mm9, mus musculus genome assembly MGSCv37; nt, nucleotide; RNA, ribonucleic acid; Stx6, syntaxin 6.

| **Gene** | **circRNA coordinates (mm9)** | **Denotation** | **circBase ID** | **circBase spliced length (nt)** |
| --- | --- | --- | --- | --- |
| N/A | chrX:58436423\|58439349 | *Cdr1as* | mmu_circ_0001878 | 2927 |
| *Stx6* | chr1:157021116\|157044565 | *Stx6 c* | mmu_circ_0000091 | 656 |
| *Ankib1* | chr5:3747021\|3772787 | *Ankib1 c* | mmu_circ_0001311 | 873 |
| *Gigyf2* | chr1:89260679\|89276583 | *Gigyf2 c* | mmu_circ_0000057 | 494 |

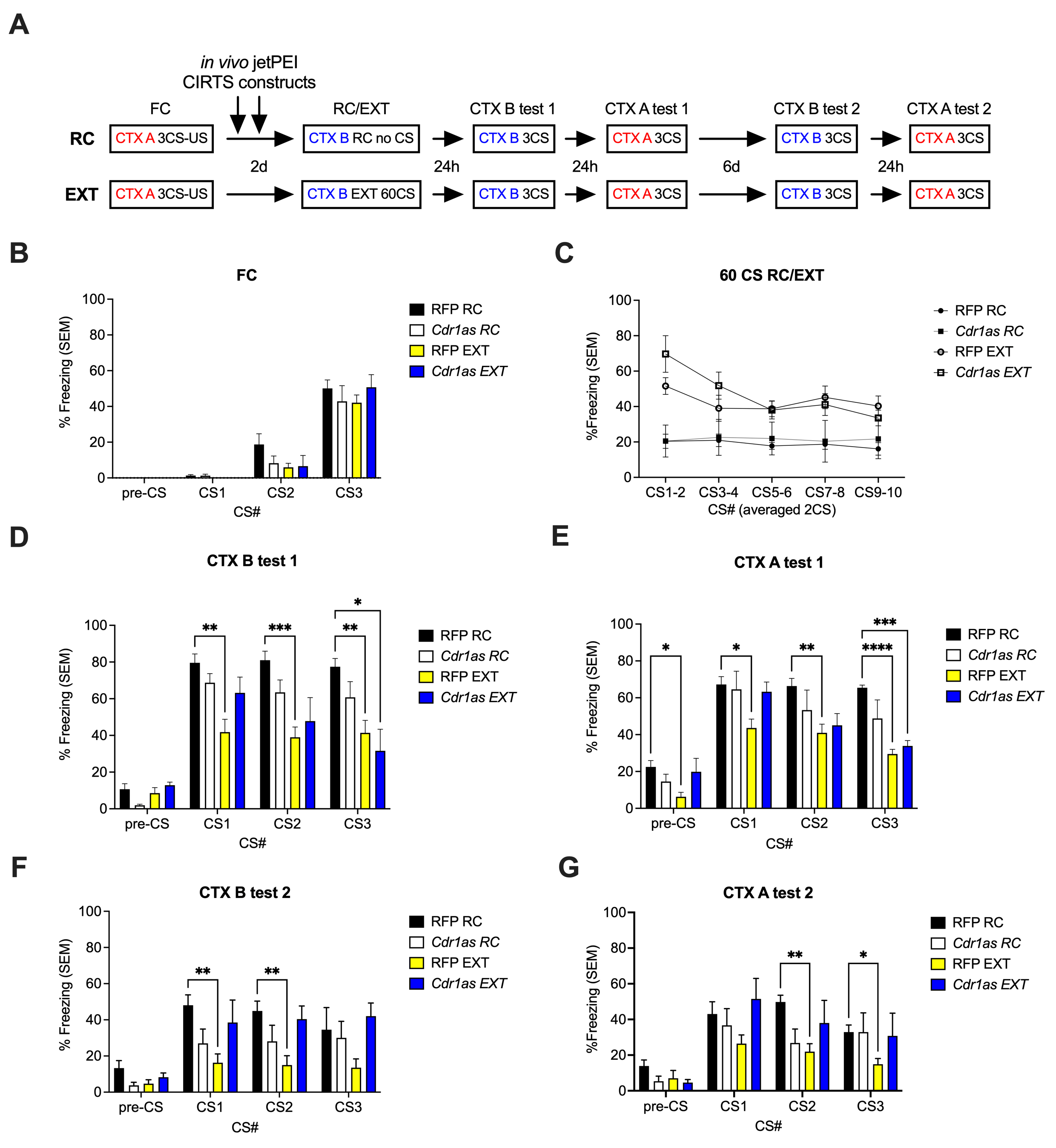

**Figure A.1:** Knockdown of Cdr1as in the ILPFC of adult male C57BL/6J mice impairs fear extinction memory. **A)** Behavioural timeline for the experiment investigating the effect of circRNA Cdr1as in fear extinction memory. Cdr1as was transiently knocked down in neurons of the ILPFC after fear conditioning and prior to extinction training using in vivo jetPEI transfection of CIRTS constructs that contain the CIRTS system with a Pin nuclease that is directed towards the neural processes via the inclusion of a Calm3 intron localisation signal (Sharangdhar et al., 2017), which is activated by neural activity, in conjunction with gRNAs antisense to the target BSJ expressed off the same construct (see Section 2.3.). **B)** All animals were fear conditioned using a 3x pairing of CS-US (tone and footshock) in context A (CTX A) and then assigned to their gRNA treatment group (RFP, Cdr1as) and behavioural group (RC, EXT) (see Section 2.4.). Two-way ANOVA (F_3,18_=1.439, P=0.2644, Dunnett’s post hoc tests relative to RFP RC control animals for all timepoints are non-significant (ns)). **C)** For RC/EXT training, animals were placed into a new context B (CTX B) and either exposed to 60x CS presentations (EXT) or left in CTX B for the same amount of time with no CS presentations (RC). **D) – G)** Context recall tests in either CTX B (D, F) or CTX A (E, G) were performed using 3x CS presentations either in the days immediately following RC/EXT training (D, E) or a week later (F, G). The amount of time they spent freezing in each context (%Freezing) is used to measure how well they recall their training; a high freezing score indicates more fear whereas a low freezing score indicates less fear. For EXT animals, it is expected that they will have less freezing compared to RC controls. RFP EXT shows significant differences to RFP RC controls. For B)-G), the two-way ANOVA results for group (outlined in figure legend) is as follows: B) F_3,18_=1.439, P=0.2644, C) F_3,21_=4.510, *P=0.0136, D) F_3,18_=6.657, *P=0.0032, E) F_3,18_=5.808, *P=0.0059, F) F_3,18_=3.919, *P=0.0258, G) F_3,18_=1.736, P=0.1954. Asterisks on the graphs indicate significance level using Dunnett’s post hoc tests relative to RFP RC control animals: *P<0.05, **P<0.01, ***P<0.001, ****P<0.0001. n = 5-7/group. All error bars depict SEM. Abbreviations: CDR1, Cerebellar Degeneration Related Protein 1; Cdr1as, CDR1 antisense RNA; c = circRNA, circular RNA; CS, conditioned stimulus; CTX, context; d, days; EXT, extinction; FC, fear conditioning; h, hours; n, number; ns, not significant; RC, retention control; RFP, red fluorescent protein; RNA, ribonucleic acid; r.t., relative to; SEM, standard error of the mean; US, unconditioned stimulus.
